# OpenMM 7: Rapid Development of High Performance Algorithms for Molecular Dynamics

**DOI:** 10.1101/091801

**Authors:** Peter Eastman, Jason Swails, John D. Chodera, Robert T. McGibbon, Yutong Zhao, Kyle A. Beauchamp, Lee-Ping Wang, Andrew C. Simmonett, Matthew P. Harrigan, Chaya D. Stern, Rafal P. Wiewiora, Bernard R. Brooks, Vijay S. Pande

## Abstract

OpenMM is a molecular dynamics simulation toolkit with a unique focus on extensibility. It allows users to easily add new features, including forces with novel functional forms, new integration algorithms, and new simulation protocols. Those features automatically work on all supported hardware types (including both CPUs and GPUs) and perform well on all of them. In many cases they require minimal coding, just a mathematical description of the desired function. They also require no modification to OpenMM itself and can be distributed independently of OpenMM. This makes it an ideal tool for researchers developing new simulation methods, and also allows those new methods to be immediately available to the larger community.

## 1. Introduction

### 1.1 Background

Molecular dynamics simulation is a rapidly advancing field. Many aspects of it are subjects of current research and development. Some of the more important examples include the development of new force fields [1,2], sometimes involving novel functional forms for the interactions [3,4]; new integration algorithms [5–7]; new sampling methods [8–10]; and support for new types of hardware [11–13].

There are many popular software packages for conducting molecular dynamics simulations. They vary considerably in their capabilities and feature sets. This is especially true when it comes to cutting edge, recently developed simulation techniques. The inventor of a new method will typically implement it in a single package, whichever one they are most comfortable working with. From that point, it may take years to be implemented in other packages, depending on the interests and priorities of the development team behind each one. In many cases, it may never get implemented. Even the initial implementation may not be accepted into an official release of the package it was created in. Or it may have limited usefulness, for example because it executes slowly or cannot be used on advanced hardware such as graphics processing units (GPUs).

The main reason for this problem is that most molecular dynamics packages were not designed with extensibility in mind. Adding new features, even very simple ones, is often labor intensive and requires a deep understanding of the code. Once a prototype implementation is complete, it may be even more difficult to turn that into a well optimized version that works on all hardware types. In most cases the simulation engines are also monolithic, so the only way to add features to them is to directly modify their source code. There is no plugin interface or other mechanism for allowing new features to be implemented and distributed independently. This turns the core development team into gatekeepers, restricting what features can be added to the package.

A complete molecular dynamics package is, of course, much more than just a simulation engine. Each one typically has its own collection of tools for preparing molecular systems to simulate, its own file formats, and sometimes even its own force fields. This makes it difficult for users to switch back and forth between them, or to combine features from different packages.

### 1.2 OpenMM

OpenMM is a molecular dynamics package designed to address these problems. It began as simply a library for performing certain types of calculations on GPUs, but in recent versions has grown into a complete simulation package with unique and powerful features. This article describes OpenMM 7.0, which is the latest release at the time of writing. An earlier version (OpenMM 4.1) was described in a previous publication [14]. This article focuses primarily on what has changed since that version, but for completeness there is some overlap between the two.

OpenMM is based on a layered architecture which (see Figure 1), to the best of our knowledge, is unique among molecular dynamics packages. This allows it to be used in several different ways by users with varying needs and interests. Depending on how a particular user chooses to interact with it, OpenMM can act as:

1. **A high-performance library**, callable from other programs, for performing a wide range of calculations used in molecular modelling and simulation on a range of advanced hardware platforms (both CPUs and GPUs).
2. **A domain specific language** for easily implementing new algorithms for molecular modelling and simulation.
3. **A complete package** for running molecular simulations.

**Figure 1:**
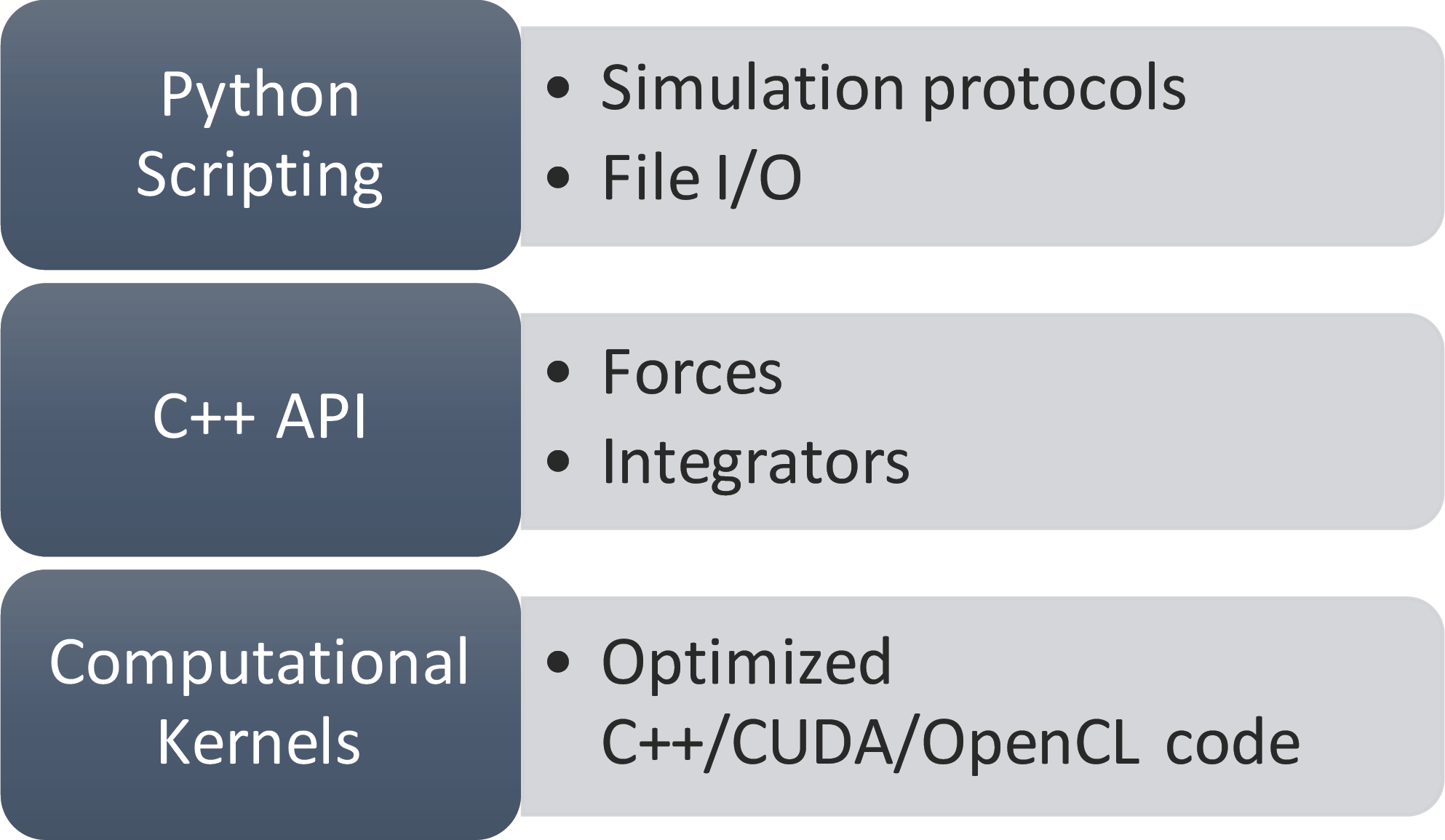
Architecture of OpenMM.

Extensibility is built into every layer of the architecture as a fundamental design goal. A guiding philosophy is that users should be able to implement new features as easily as possible, by writing as little code as possible. Those features should then work on all types of hardware, including both CPUs and GPUs, and have good performance on all of them. Finally, the developer of a feature should be able to package and distribute their code independently, without needing the approval or participation of the core OpenMM development team.

The highest layer of the architecture is based on the Python scripting language. Users can easily extend it by writing their own Python code to implement the algorithms of their choice. A wide range of simulation protocols, sampling methods, etc. can be implemented in this way, often with only a few lines of code.

The next layer down defines the calculations that are tied together through Python scripting. This layer includes many classes for creating “custom” forces and integrators. These classes provide a simple but powerful mechanism for extensibility. The user provides one or more mathematical expressions to describe the calculation to be done. For example, they might give an expression for the interaction energy of a pair of particles as a function of the distance between them. The expression is parsed and analyzed, and just-in-time compilation is used to generate an efficient implementation of the code for calculating that interaction [15]. This allows users to easily define a huge variety of interactions and integration algorithms. They can then be used on any supported type of hardware, and involve little or no loss in performance.

At the lowest layer, OpenMM is based on a plugin mechanism. Calculations are defined by “computational kernels”. A plugin may define new kernels for doing new types of calculations, or alternatively it may provide new implementations of existing kernels, for example to support a new type of hardware. Plugins are dynamically discovered and loaded at runtime. Each one is packaged as a file that can be distributed separately from the rest of OpenMM and installed by any user.

Another unique feature of OpenMM is its support for multiple input pipelines. Before a molecular system can be simulated, it first must be modelled. This is sometimes a complex process involving such steps as combining multiple molecules into a single file, building missing loops, selecting a force field, and parametrizing small molecules. Typically, each simulation package provides its own tools for doing this. They often differ in significant ways, such as what force fields are available.

OpenMM does include modelling tools, but it also can directly read the file formats used by Amber [16], CHARMM [17], Gromacs [18], and Desmond [19]. A user can prepare their system with the tools from any of those packages, or with other tools that are designed to work with them, then simulate it in OpenMM. This gives great flexibility, since the user can use whatever tools are best suited to the system they want to simulate. It also lets OpenMM easily fit into their existing workflow. A user who is accustomed to a particular tool can continue to use it, but still run their simulation in OpenMM.

## 2. Design and Implementation

### 2.1 Extensibility

#### Python Scripting

The highest level of the architecture consists of a set of Python classes and functions. They may be chained together to create simple scripts that run simulations, or more complicated ones that implement a variety of advanced algorithms. These are some of the functions provided by OpenMM that may be used by Python scripts:

- Reading input files, including standard formats like PDB or PDBx/mmCIF, as well as the proprietary formats used by applications such as Amber, CHARMM, Gromacs, and Desmond.
- Editing molecular models, such as by combining molecules together, adding or deleting atoms, building solvent boxes, etc.
- Defining the forces acting on a molecular system, either by specifying them explicitly or by loading a force field definition from a file.
- Computing forces and energies.
- Running simulations.
- Outputting results.

An example of a script to run a simulation is shown in Listing 1. It loads a PDB file, models the forces with the AMBER99SB-ILDN force field [20] and TIP-3P water model [21], performs a local energy minimization to eliminate clashes, and then simulates 1 million steps of Verlet dynamics. Every 1000 steps, it writes the current structure to a DCD file, and the current time, potential energy, and temperature to a log file.

**Listing 1:**
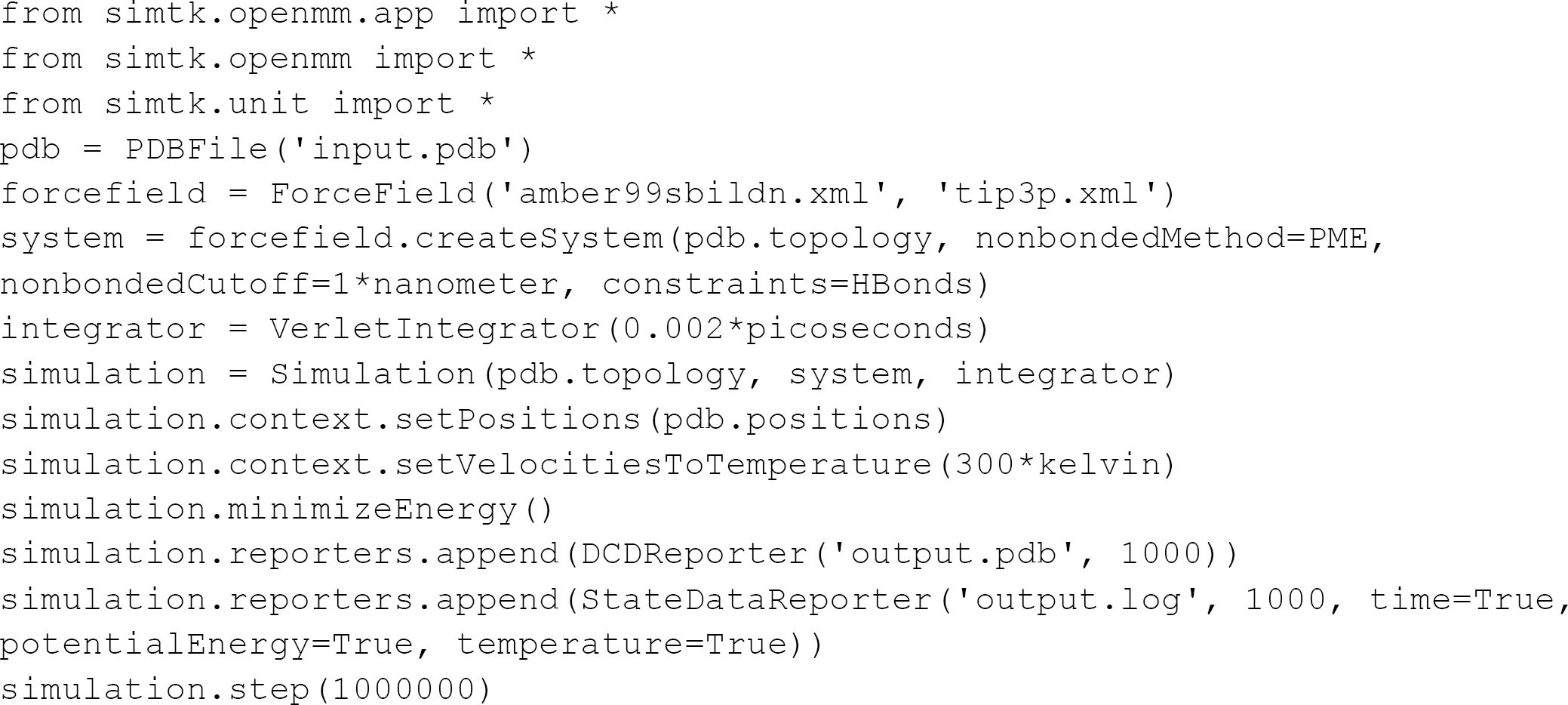
A Python script executing a complete molecular simulation from a PDB file.

This script runs a simple simulation, much like one might run in any molecular dynamics package. More sophisticated or exotic algorithms and protocols can be implemented in exactly the same way. Here are some examples of features or applications that use or extend OpenMM’s Python scripting features:

- A class for simulated tempering [10], an accelerated sampling method that varies the temperature of a simulation to accelerate barrier crossings. The entire algorithm was implemented in roughly 200 lines of code.
- PDBFixer, an application for cleaning up molecular models in preparation for simulating them. It includes such features as building missing loops, replacing nonstandard amino acids with standard ones, adding hydrogens, and building solvent boxes. By using the features provided by OpenMM, all of these algorithms were implemented in only about 1000 lines of code.
- YANK [22] a sophisticated application and toolkit for alchemical free energy calculations. It implements Hamiltonian exchange molecular dynamics simulations to efficiently sample multiple alchemical states, and utilizes the “custom” forces provided by OpenMM to allow exploration of many different alchemical intermediate functional forms.

Although these tools are written in Python, all expensive calculations are done by OpenMM and take full advantage of the available hardware, including GPUs and multicore CPUs. Because they interact with OpenMM only through well-defined public interfaces, they can be packaged and distributed independently. No changes to OpenMM itself are required to use them.

A set of advanced examples is included in the supporting information. They demonstrate more complex simulation techniques, and show how to use OpenMM in combination with other programs.

#### Custom Forces

In addition to the standard forces provided by OpenMM (such as Lennard-Jones forces, PME and reaction field electrostatics, and generalized Born models), custom forces are a mechanism for creating interactions between particles with entirely novel functional forms. There are many different custom force classes, each supporting a particular category of interactions. They are listed in Table 1. Since OpenMM 4.1 was described in an earlier publication, several new custom force classes have been added, including CustomCompoundBondForce (added in OpenMM 5.0), CustomManyParticleForce (added in OpenMM 6.2), and CustomCentroidBondForce (added in OpenMM 7.0).

**Table 1:** **Custom forces supported by OpenMM 7.0**.

| Custom Force Class | Description |
| --- | --- |
| CustomBondForce | Applies forces to pairs of bonded atoms based on the distance between them. |
| CustomAngleForce | Applies forces to triplets of bonded atoms based on the angle between them. |
| CustomTorsionForce | Applies forces to sets of four bonded atoms based on the dihedral between them. |
| CustomExternalForce | Applies forces to individual atoms based on their positions. |
| CustomCompoundBondForce | Applies forces to sets of arbitrarily many bonded atoms based on any combination of their positions, distances, angles, and dihedrals. |
| CustomNonbondedForce | Applies forces to pairs of non-bonded atoms based on the distance between them. |
| CustomGBForce | Supports multi-stage computations of non-bonded interactions, such as generalized Born implicit solvent models. |
| CustomCentroidBondForce | Similar to CustomCompoundBondForce, but the interaction is based on the centroids of groups of atoms rather than individual atoms. |
| CustomManyParticleForce | Supports non-bonded interactions that depend on the positions of arbitrarily many atoms at once. |
| CustomHbondForce | Supports a variety of hydrogen bonding models. |

In each case, the user provides an algebraic expression for the interaction energy as a function of the relevant variables. OpenMM analytically differentiates the expression to determine the corresponding force, then uses just-in-time compilation to generate machine code for efficiently computing the force and energy on the current hardware (either CPU or GPU).

As an example, Listing 2 defines a harmonic restraint that can be applied to the angles formed by triplets of atoms. It specifies that the energy of each triplet is given by *k*(*θ*-*θ*_0_)^2^. It also specifies that *k* and *θ*_0_ are per-angle parameters: each triplet can have different values for them.

**Listing 2:**
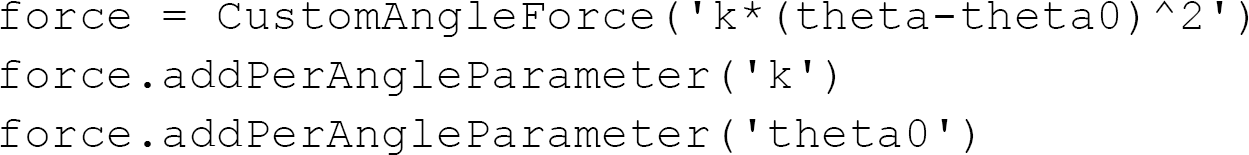
Implementation of a harmonic angle restraint using a CustomAngleForce.

Custom forces are designed to achieve several goals that usually conflict with each other. First, it should be exceptionally easy to implement completely new functional forms for interactions. As seen in Listing 2, it often requires no more than a few lines of Python code. Second, a single implementation should work on all types of hardware. The exact same code can be used whether the program is being run on a CPU or GPU. Third, the user should not need to sacrifice performance. Because the expression is converted to machine code before the simulation is run, there often is little or no difference in speed between a custom force and a hand-written implementation of the same interaction.

#### Custom Integrators

Just as custom forces allow users to implement novel interactions, custom integrators allow them to implement novel integration algorithms. The algorithm is defined by a sequence of operations, each defining a calculation to be done. Various types of operations are supported. Examples include:

- Evaluating a mathematical expression for each degree of freedom, then assigning the result to a variable for each one.
- Evaluating a mathematical expression once and assigning the result to a global variable.
- Summing an expression over all degrees of freedom and assigning the result to a global variable.
- Applying constraints to positions or velocities.

In the simplest case, all operations are executed in order to take a single integration time step. In addition, OpenMM 7.0 added support for more complex flow control through *if* and *while* blocks.

Listing 3 shows Python code that uses a custom integrator to implement the leapfrog Verlet algorithm. The function `addPerDofVariable()` defines a new variable that has a different value for each degree of freedom. The function `addComputePerDof()` defines a calculation to be performed independently for each degree of freedom. In the absence of constraints, each time step of this algorithm consists of the operations

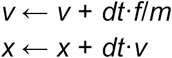

where *x* is the position at time *t*, *v* is the velocity at time *t*-*dt*/2, *dt* is the step size, *f* is the force, and *m* is the particle mass. When constraints are present, the positions must then be adjusted to satisfy them, and finally the velocities are recalculated as

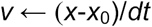

where *x*_0_ is the position at the start of the step.

**Listing 3:**
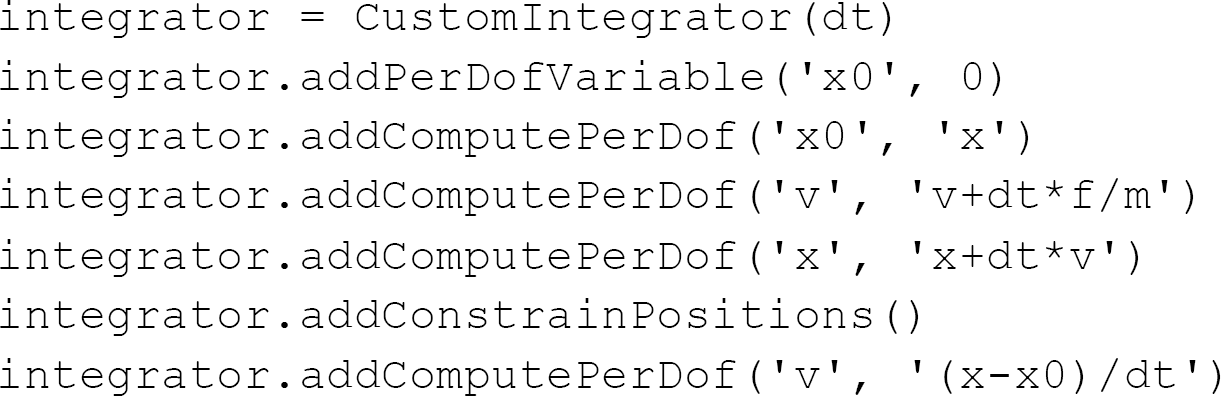
Leapfrog Verlet algorithm implemented as a CustomIntegrator.

Far more complicated and sophisticated algorithms can be implemented in the same way. Here are some examples of integrators that have been created with this mechanism.

- The rRESPA multiple time step integration algorithm [23].
- The aMD accelerated sampling algorithm [9].
- Metropolis-Hastings Monte Carlo [24] with Gaussian displacement proposals.
- Hybrid Monte Carlo and variants, such as Generalized hybrid Monte Carlo (GHMC) [25], a Metropolized form of Langevin dynamics.
- Nonequilibrium candidate Monte Carlo (NCMC) [26], where an external field is changed during the course of dynamics and the resulting nonequilibrium proposal accepted or rejected to preserve the equilibrium distribution.

As with custom forces, a single implementation works on all types of hardware. Because just-in-time compilation is used to generate efficient machine code for the algorithm, there usually is little or no performance cost relative to using handwritten GPU code.

#### Plugins

The lowest layer of the OpenMM architecture is based around plugins. This allows new features to be packaged as libraries, distributed independently, and loaded dynamically at runtime. For example, a plugin can implement a new type of interaction or a new integration algorithm, or it can add support for a new type of hardware. In fact, many of the core features of OpenMM are actually implemented as plugins, including its implementations of the AMOEBA force field [27], ring polymer molecular dynamics (RPMD) [28], and polarizable Drude particles [29].

This provides nearly unlimited extensibility, allowing users to implement any feature they might want. Writing a plugin involves far more work than the other extensibility features described above. For example, it is up to the user to write whatever code is necessary to make it work on each type of hardware, such as CUDA or OpenCL code for GPUs. When possible, it is therefore usually preferable to use one of the other mechanisms. Nevertheless, plugins are an important option when extreme extensibility and performance is needed. Like the other mechanisms, they allow a developer to create an extension and distribute it directly to users. No modifications to OpenMM itself are needed.

### 2.2 Advanced Features

OpenMM has many other features beyond those discussed above, some of which are themselves unique or noteworthy. The following are some of the more significant ones.

#### AMOEBA

OpenMM has an implementation of the AMOEBA polarizable force field which is, to the best of our knowledge, the fastest available in any code [3]. AMOEBA is designed to transcend the limitations of conventional point charge force fields and achieve much higher accuracy in force and energy computations. It uses two main mechanisms to achieve this. First, instead of approximating atoms as point charges, it assigns each one a multipole moment up to the level of quadrupoles. Second, it explicitly models atomic polarization by assigning an induced dipole to each atom. Because the induced dipoles interact with each other, they must be computed at each time step using an iterative self-consistent field calculation. Both of these features make AMOEBA far slower to simulate than conventional force fields.

Much research has been done recently on ways to reduce this cost, and new versions of OpenMM have incorporated several of the most recent algorithms. Interactions between multipoles are computed using spherical harmonics in a quasi-internal coordinate system [30,31] (added in OpenMM 7.0). The iterative solver for induced dipoles uses the Direct Inversion in the Iterative Subspace (DIIS) algorithm [32] (added in OpenMM 6.1). Alternatively, it can use the recently developed extrapolated polarization approximation [33] (added in OpenMM 7.0). In this method, only a few iterations are performed, and then an analytic approximation is used to extrapolate to the limit of infinite iterations. This can give a large improvement in speed with only a very small loss in accuracy.

#### Drude Particles

A new feature introduced in OpenMM 5.2 is support for Drude particles [29] as an alternative way of modelling polarizability. In this method, each polarizable atom is modelled as a pair of charges connected by an anisotropic harmonic force. When an electric field is applied, the two particles are displaced from each other, creating a dipole moment. The strength of the force connecting them determines the atomic polarizability. The particle positions can be determined using a self-consistent field calculation or, more commonly, a dual-thermostat Langevin integrator that couples the center of mass of each pair to a high temperature heat bath (e.g. 300K), but the internal motion of each pair to a low temperature heat bath (e.g. 1K).

Polarizable force fields based on Drude particles are included with OpenMM. This includes the SWM4-NDP water model [34], and the CHARMM polarizable force field for proteins [4]. They aim to incorporate some of the same physical effects as AMOEBA at a lower computational cost.

#### Virtual Sites

Virtual sites are interaction sites within a molecule whose positions are not integrated directly. Instead, they are calculated at each time step based on the positions of other particles. They are often used to provide a more detailed charge distribution than would be possible using only a single point charge for each atom. For example, they appear in many multisite water models (such as TIP-4P and TIP-5P), and also in the CHARMM polarizable protein force field.

There are a multitude of possible ways a virtual site position can be specified based on the positions of other mobile atoms. Typically, a simulation package will provide a limited choice of rules, covering only those cases needed for the particular force fields that package supports. For example, Gromacs 5 offers a choice of four methods for calculating a virtual site position based on the positions of three atoms. Each one covers one very specific case that occurs in a supported force field. One can easily imagine other cases that would be impossible to construct with any of the current rules, and would therefore require adding a fifth rule.

OpenMM also offers a few specialized rules for positioning virtual sites, but in addition, it has a very general method designed to cover all cases that are ever likely to occur in which a virtual site depends on the positions of three atoms. This method, added in OpenMM 6.1, can reproduce all four of the rules provided by Gromacs, as well as supporting many other situations they could not.

In this method, three vectors are first calculated as weighted averages of the positions of the three atoms:

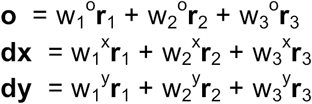

where **r**_1_, **r**_2_, and **r**_3_ are the atom positions, and the coefficients are user-defined. They are then used to construct a set of orthonormal coordinate axes 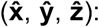

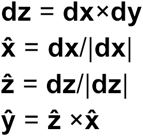

Finally, the virtual site position is set to an arbitrary user-defined location within this coordinate system:

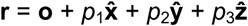

This method is another example of how flexibility and extensibility are core design goals of OpenMM. Instead of supporting only a limited set of specialized virtual site types, it tries to provide a very general type that can cover as wide a range of cases as possible, thus giving maximum flexibility to users in designing their models and force fields.

#### Triclinic Periodic Boxes

Earlier versions of OpenMM supported only rectangular periodic boxes. In OpenMM 6.3 it was extended to support triclinic boxes as well: ones formed by combinations of three arbitrary lattice vectors. It can be shown that this formulation is extremely general; all standard periodic box shapes, including the popular rhombic dodecahedron and truncated octahedron, can be represented as triclinic boxes [35].

This feature serves two important functions. First, it allows one to simulate crystals, which very often have non-rectangular unit cells. Second, it is useful when simulating freely rotating molecules in solvent. One needs to include a certain amount of padding around the molecule to ensure that no two periodic copies ever come too close together. Because the molecule can freely rotate, the same padding is required along all directions, so one wants the periodic box to be as close as possible to spherical. For a given padding distance, the rhombic dodecahedron has only about 71% the volume of a rectangular unit cell. It therefore requires less solvent and reduces the cost of the simulation.

#### Multiple Precision Modes

Many aspects of a molecular dynamics code involve tradeoffs between speed and accuracy. This is especially true when executing on a GPU, since they often have very poor double precision performance. To optimize execution speed, it is preferable to use single precision whenever possible, resorting to double precision only when absolutely necessary. Unfortunately, there is no unique standard for when it is “necessary”. The minimum acceptable level of error can vary widely depending on the details of a simulation and the type of information one wishes to obtain from it.

OpenMM 5.0 introduced support for multiple precision modes. When running on a GPU, the user has a choice of three modes:

- *Single Precision:* Nearly all calculations are done in single precision. Double is used in only a handful of places where it has negligible impact on performance and is most important for accuracy.
- *Mixed Precision:* Forces are computed in single precision, but integration and energy accumulation are done in double precision. This gives a large improvement in the accuracy of some quantities, while only having a small impact in performance.
- *Double Precision:* All calculations are done in double precision. This gives the best accuracy, but often has a very large effect on performance.

The effects of the different modes are illustrated below in Section 3.

Regardless of the precision mode, forces are accumulated as 64 bit fixed point values. This improves accuracy when working in single or mixed precision modes, and ensures that force accumulation is deterministic. It also allows force accumulation to be done with integer atomic operations, which substantially improves performance. OpenMM has used this method since version 4.0, released in January 2012. Since that time, it has found its way into other GPU accelerated MD codes, such as AMBER [36].

## 3. Results

### 3.1 Performance

To evaluate the speed of OpenMM, we benchmarked its performance with three molecular systems of varying size:

1. Dihydrofolate reductase (DHFR), a 2489 atom protein solvated with 7023 water molecules to give a total of 23,558 atoms.

2. Abl kinase (ABL1), a 4067 atom protein solvated with 13,692 water molecules to give a total of 45,143 atoms

3. The mechanistic target of rapamycin (MTOR), a 19,019 atom protein solvated with 56,733 water molecules to give a total of 189,218 atoms.

Benchmarks were run on the following types of hardware:

1. An NVIDIA Titan X Pascal GPU.

2. An NVIDIA Tesla K80 GPU.

3. A 4 core, 3.5 GHz Intel Core i7-2700K CPU.

All GPU simulations used CUDA 7.5. The K80 consists of two independent GPUs on a single board. OpenMM can parallelize a single simulation across multiple GPUs, or alternatively run a different simulation on each one at the same time. We therefore included benchmarks using only one of the GPUs (thus leaving the other free for a different simulation), as well as ones using both GPUs for a single simulation.

In the discussion below, we summarize the most important parameters for each set of simulations. Full details can be found in the scripts used to run the simulations, which are included in the Supplemental Information.

#### AMBER

We first benchmarked the performance using the AMBER99SB-ILDN force field and TIP3P water model. All simulations used a Langevin integrator with a temperature of 300 K and a friction coefficient of 1 ps^-1^. Long range Coulomb interactions were computed with the Particle Mesh Ewald (PME) method.

Simulations were run using integration time steps of both 2 fs and 5 fs. For the 2 fs simulations, covalent bonds involving a hydrogen atom were modelled as rigid constraints. For the 5 fs simulations, all covalent bonds were modelled as rigid constraints and hydrogen mass repartitioning was used to increase the mass of hydrogen atoms to 4 amu (while decreasing the masses of the atoms they were bonded to so as to keep the total system mass constant). In all cases, water molecules were kept rigid. All of the GPU simulations used single precision.

The results are shown in Table 2.

**Table 2:**
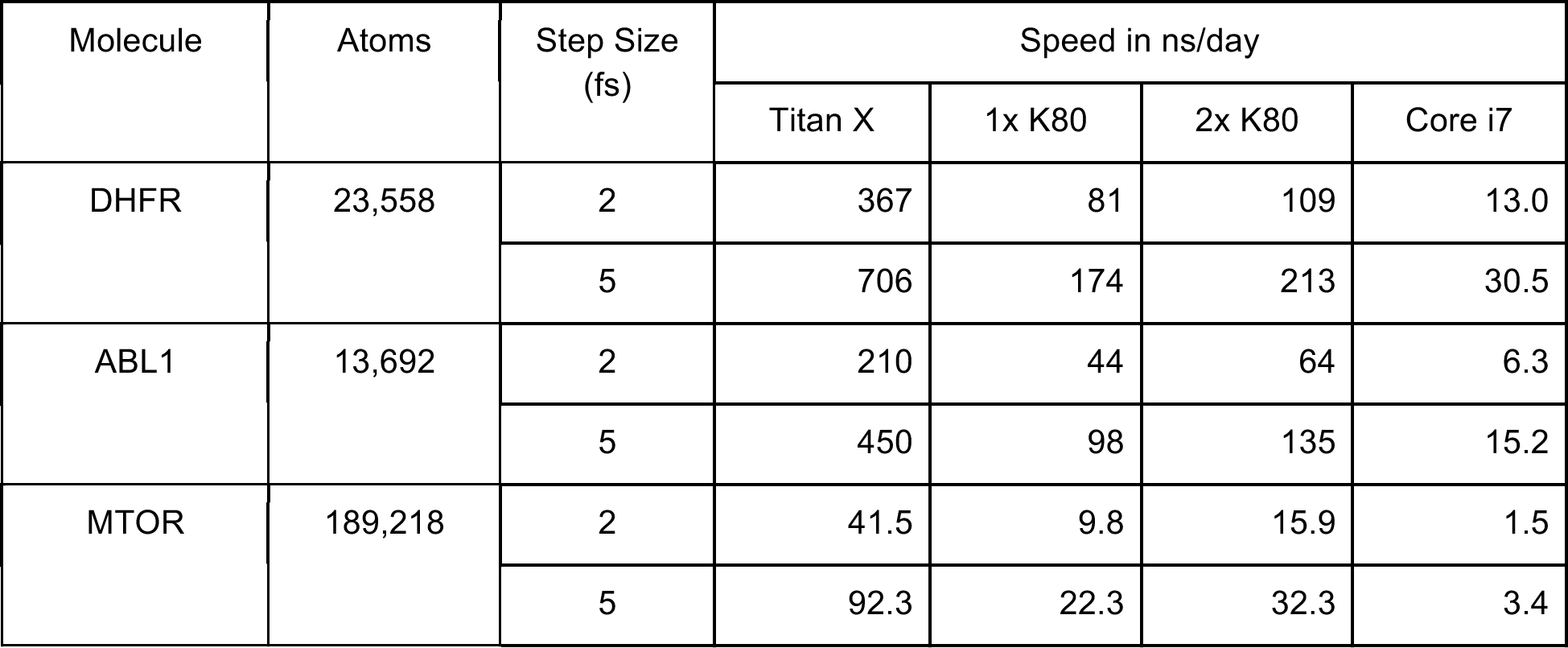
Benchmark results for various protein systems in explicit solvent simulated with PME.

Depending on the molecule and settings, using two GPUs is anywhere from 22% to 62% faster than a single GPU, with the larger molecules generally having the higher speedups. If the goal is to run a single simulation as quickly as possible, using multiple GPUs is therefore quite useful. On the other hand, if the goal is to generate as much total simulation time as possible, it is more efficient to run a separate simulation on each one.

#### AMOEBA

We next benchmarked performance using the AMOEBA2013 force field. The AMOEBA water model is designed to be flexible rather than rigid, which requires a smaller step size. We therefore used a rRESPA multiple time step integrator, in which bonded forces were evaluated every 1 fs and nonbonded forces every 2 fs. No degrees of freedom were constrained. As above, we used PME for long range Coulomb interactions and single precision.

Simulations were run with two different methods of calculating the induced dipoles:

1. Full mutual polarization with a tolerance of 10^-5^ for the induced dipoles.

2. The extrapolated polarization approximation.

The CPU implementation of AMOEBA in OpenMM is not well optimized, so we only ran benchmarks on GPUs. Using multiple GPUs for a single simulation is not supported with AMOEBA. Because AMOEBA is a very expensive force field and is normally only used for modest sized systems, we only ran benchmarks for DHFR.

The results are shown in Table 3.

**Table 3:** Benchmark results for DHFR in explicit solvent using AMOEBA2013.

| Polarization | Speed in ns/day |  |
| --- | --- | --- |
|  | Titan X | K80 |
| Mutual | 10.09 | 2.84 |
| Extrapolated | 20.90 | 4.58 |

#### Effect of Precision

When running on a GPU, OpenMM gives a choice of three precision modes: single, mixed, and double. To measure the effect of this choice on performance, we repeated the 2 fs time step DHFR simulations in mixed and double modes. The results are shown in Table 4.

**Table 4:** Effect of precision model on performance.

| Precision | Speed in ns/day |  |
| --- | --- | --- |
|  | Titan X | K80 |
| Single | 367 | 81 |
| Mixed | 332 | 78 |
| Double | 18.1 | 30.2 |

The speed difference between single and mixed precision is quite small, whereas double precision is much slower. This is especially true on the Titan X, a GPU primarily targeted at consumers that has very poor double precision performance. The Tesla K80, which is targeted at high performance computing, does much better, although there is still a large decrease in performance. Overall, the Titan X is far faster in single or mixed precision modes, while the K80 is faster in double precision mode.

To see the benefits of higher precision, we performed additional simulations of DHFR. Because a thermostat tends to mask the effect of error, these simulations used a leapfrog Verlet integrator to simulate constant energy. All simulations used the AMBER99SB-ILDN force field, a 2 fs time step, rigid water, and constraints on bonds involving hydrogen.

Each simulation was 1 ns in length. The total energy was recorded every 1 ps, and a linear regression was used to estimate the rate of energy change. Ten independent simulations were performed for each precision mode, giving ten estimates of the rate. Table 5 reports the mean and standard error of those ten rates for each mode.

**Table 5.** Energy drift for different precision models for the DHFR explicit solvent system. The reported uncertainty in each value is the standard error of the drift rates from ten independent simulations.

| Precision | Energy drift in (kJ/mol)/ps |
| --- | --- |
| Single | 1.557 ± 0.003 |
| Mixed | -0.0047 ± 0.0008 |
| Double | -0.0062 ± 0.0002 |

The energy drift in single precision is more than two orders of magnitude larger than in mixed or double precision. When accurate energy conservation is important, using mixed precision has a very large benefit at low cost. The average drift rates in mixed and double precision are not significantly different from each other, indicating that numeric precision is no longer the dominant source of error. In other cases, such as when using a smaller step size or when simulating a larger molecule, statistically significant differences between them might emerge.

### 3.2 Input Pipelines

A key feature of OpenMM is its support for multiple input pipelines. This allows users to prepare molecular systems with the tools of their choice, then simulate them in OpenMM. Support for Gromacs input files was added in OpenMM 5.1, Desmond file support was added in OpenMM 6.0, and CHARMM file support was added in OpenMM 6.1.

The code in Listing 1 began from a PDB file and force field definition, using those to construct a description of the molecular system. Listing 4 shows the changes needed to instead construct it from an Amber prmtop file, as created by the AmberTools suite of software. More complete examples of using Amber and CHARMM input files are included in the supporting information.

**Listing 4:**
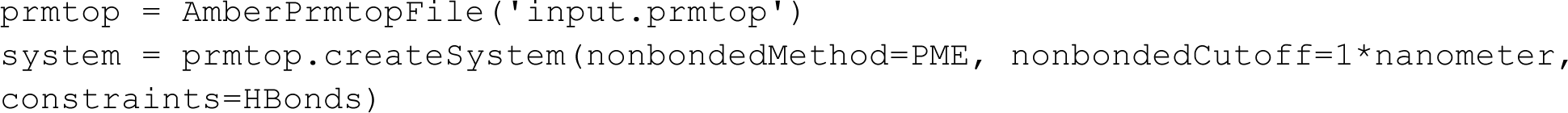
Loading a system from Amber prmtop/inpcrd files.

To validate the accuracy of the input pipelines, we constructed systems using the setup tools from other packages, then loaded them into those packages and into OpenMM and compared the forces and energies. We performed these tests on two systems: DHFR, a 159 residue protein, and 2KOC, a 14-mer hairpin RNA. Comparisons were made to Amber 16, Gromacs 4.6.5, and CHARMM-LITE c40b1. For Amber, we performed comparisons in both explicit solvent and OBC1 implicit solvent. For Gromacs and CHARMM, we compared only explicit solvent.

To create the Amber input files, ParmEd [37] was used to download PDB files 4M6J (DHFR) and 2KOC (RNA hairpin), which were then stripped of water molecules and adjusted to standard amino acids. In the case of 2KOC, the first model was used. Cofactors and phosphates were deleted. Amber prmtop and inpcrd files were created with LEaP from the AmberTools 16 distribution. For implicit solvent simulations, mbondi3 GB radii were used; for explicit solvent simulations, the system was solvated with TIP3P waters in an octahedral box with 15 Å of clearance, and 20 Na+ and 20 Cl- counterions were added.

To create the Gromacs input files, ParmEd was used to convert the Amber prmtop and inpcrd files into Gromacs top and gro files.

To create the CHARMM input files, CHARMM-GUI [38] was used to download the 4M6J and 2KOC PDB files. Crystallographic water molecules were deleted, and the system was solvated with a rectangular water box with 15 Å of padding. Default values were accepted for all other options, including replacing nonstandard amino acids and patching terminal residues.

All of the input files, as well as scripts needed to run the comparisons, are included in the Supplemental Information.

Results are shown for Amber in Tables 6 and 7, for Gromacs in Tables 8 and 9, and for CHARMM in Tables 10 and 11. In all cases the agreement is excellent, with all energy components matching to at least four significant digits. In systems that use PME, the nonbonded energies have somewhat larger differences than other energy components. This is partly because of the larger magnitude of this interaction, and partly because of the fact that different applications compute nonbonded interactions in slightly different ways. For example, Amber uses 4th order splines for charge spreading, while OpenMM uses 5th order splines. Nonetheless, they both compute the forces and energy to similar overall accuracy.

**Table 6.** Comparison of energy components, as calculated by Amber and OpenMM. All values are in kJ/mol.

|  | 2KOC, OBC |  | 2KOC, PME |  | DHFR, OBC |  | DHFR, PME |  |
| --- | --- | --- | --- | --- | --- | --- | --- | --- |
| Term | Amber | OpenMM | Amber | OpenMM | Amber | OpenMM | Amber | OpenMM |
| Bond | 7876.38 | 7876.38 | 7877.63 | 7877.63 | 611.05 | 611.05 | 613.34 | 613.34 |
| Angle | 274.19 | 274.19 | 274.19 | 274.19 | 1611.89 | 1611.89 | 1611.89 | 1611.89 |
| Dihedral | 1416.68 | 1416.68 | 1416.68 | 1416.68 | 8844.32 | 8844.32 | 8844.32 | 8844.32 |
| Nonbonded | -3316.70 | -3316.81 | -235740.76 | -235750.52 | -21806.02 | -21806.70 | -433365.84 | -433410.31 |
| OBC | -11607.17 | -11607.57 |  |  | -13766.56 | -13767.04 |  |  |
| Total | -5356.62 | -5357.13 | -226172.26 | -226182.02 | -24505.32 | -24506.48 | -422296.30 | -422340.77 |

**Table 7.** Comparison of forces as computed by Amber and OpenMM. Values are the normalized projection of the Amber forces (**F**_A_) onto the OpenMM forces (**F**_O_): (**F**_A_·**F**_O_)/(**F**_O_·**F**_O_). The mean, minimum, and maximum are taken over all atoms.

|  | 2KOC, OBC | 2KOC, PME | DHFR, OBC | DHFR, PME |
| --- | --- | --- | --- | --- |
| Mean | 0.99999 | 0.99996 | 1.00000 | 0.99997 |
| Minimum | 0.99987 | 0.99149 | 0.99981 | 0.96374 |
| Maximum | 1.00012 | 1.00530 | 1.00021 | 1.00997 |

**Table 8.** Comparison of energy components, as calculated by Gromacs and OpenMM. All values are in kJ/mol.

|  | 2KOC |  | DHFR |  |
| --- | --- | --- | --- | --- |
| Term | Gromacs | OpenMM | Gromacs | OpenMM |
| Bond | 7976.96 | 7976.95 | 682.27 | 682.27 |
| Angle | 277.21 | 277.21 | 1646.32 | 1646.32 |
| Dihedral | 1416.77 | 1416.76 | 8847.34 | 8847.38 |
| Nonbonded | -235817.06 | -235793.81 | -433422.38 | -433449.40 |
| Total | -226146.12 | -226122.89 | -422246.45 | -422273.43 |

**Table 9.** Comparison of forces as computed by Gromacs and OpenMM. Values are the normalized projection of the Gromacs forces (**F**_G_) onto the OpenMM forces (**F**_O_): (**F**_G_·**F**_O_)/(**F**_O_·**F**_O_). The mean, minimum, and maximum are taken over all atoms.

|  | 2KOC | DHFR |
| --- | --- | --- |
| Mean | 1.00000 | 1.00000 |
| Minimum | 0.99807 | 0.99785 |
| Maximum | 1.00041 | 1.00230 |

**Table 10.**
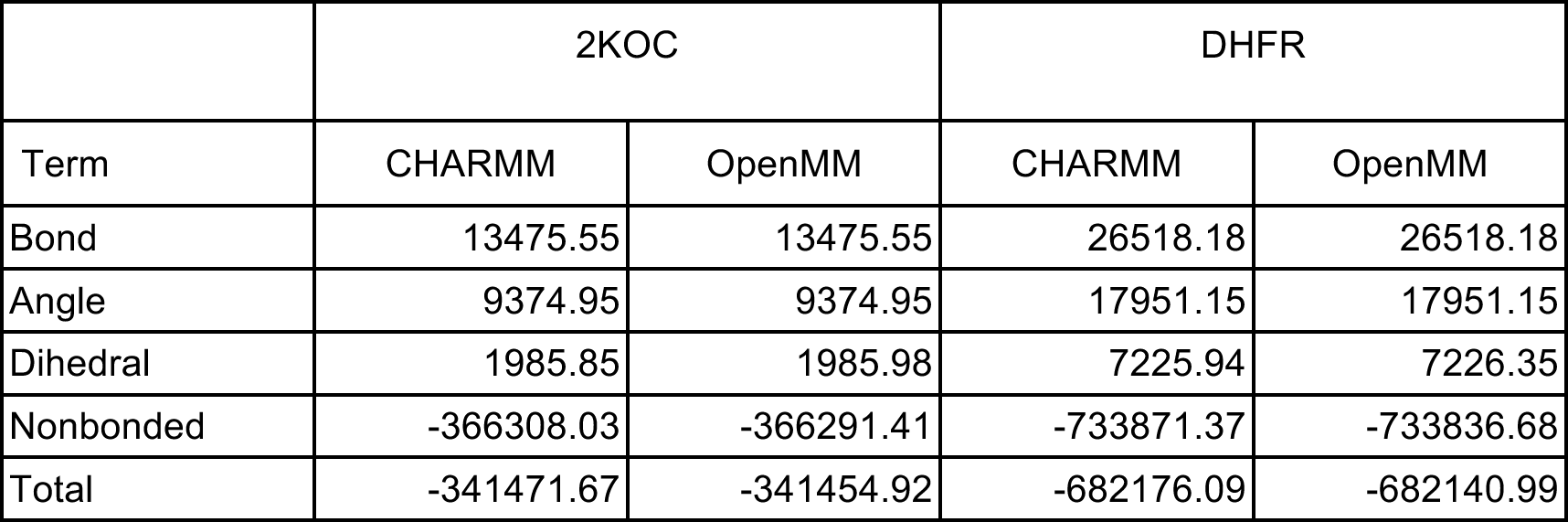
Comparison of energy components, as calculated by CHARMM and OpenMM using the CHARMM36 force field. All values are in kJ/mol.

|  | 2KOC |  | DHFR |  |
| --- | --- | --- | --- | --- |
| Term | CHARMM | OpenMM | CHARMM | OpenMM |
| Bond | 13475.55 | 13475.55 | 26518.18 | 26518.18 |
| Angle | 9374.95 | 9374.95 | 17951.15 | 17951.15 |
| Dihedral | 1985.85 | 1985.98 | 7225.94 | 7226.35 |
| Nonbonded | -366308.03 | -366291.41 | -733871.37 | -733836.68 |
| Total | -341471.67 | -341454.92 | -682176.09 | -682140.99 |

**Table 11.**
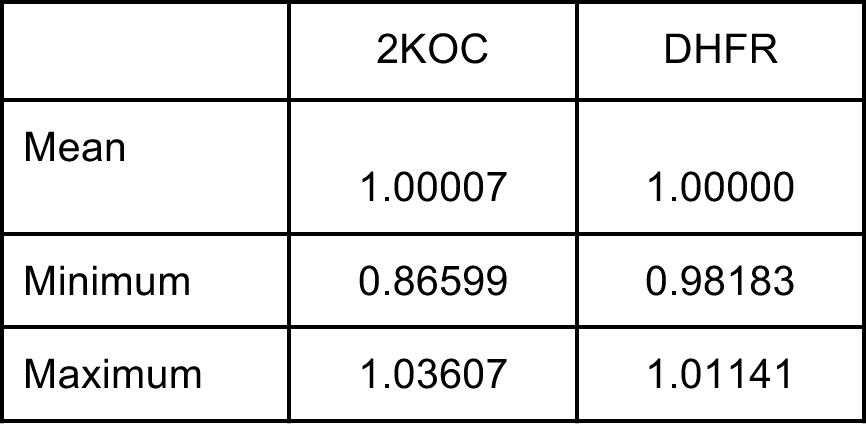
Comparison of forces as computed by CHARMM and OpenMM. Values are the normalized projection of the CHARMM forces (**F**_C_) onto the OpenMM forces (**F**_O_): (**F**_C_·**F**_O_)/(**F**_O_·**F**_O_). The mean, minimum, and maximum are taken over all atoms.

## 4. Availability and Future Directions

OpenMM is available from http://openmm.org. Ongoing development is conducted through the Github community at https://github.com/pandegroup/openmm. Detailed instructions on how to compile it from source are found in the OpenMM User Guide at http://openmm.org/documentation.html.

## 5. Acknowledgements

The authors thank Daniel L. Parton (MSKCC) for providing input files for the ABL1 and MTOR benchmarks.

## Supporting Information Legends

S1 Supporting Information. Scripts and data files to reproduce the results described in section 3.

S2 Source Code. Source code and documentation for OpenMM 7.0.1.

S3 Examples. Examples and tutorials demonstrating more advanced usage of OpenMM.

